# Context dependency of biotic interactions and its relation to plant rarity

**DOI:** 10.1101/791269

**Authors:** Anne Kempel, Hugo Vincent, Daniel Prati, Markus Fischer

## Abstract

**Aim:** Biotic interactions can determine rarity and commonness of species, however evidence that rare and common species respond differently to biotic stress is scarce. This is because biotic interactions are notoriously context-dependent and traits leading to success in one habitat might be costly or unimportant in another. We aim to identify plant characteristics that are related to biotic interactions and may drive patterns of rarity and commonness, taking environmental context into account.

**Location:** Switzerland

**Methods:** In a multi-species experiment, we compared the response to biotic interactions of 19 rare and 21 widespread congeneric plant species in Switzerland, while also accounting for variation in environmental conditions of the species’ origin.

**Results:** Our results restrict the long-standing hypothesis that widespread species are superior competitors to rare species to only those species originating from resource rich habitats, in which competition is usually strong. Tolerance to herbivory and ambient herbivore damage on the other hand, did not differ between widespread and rare species. In accordance to the resource-availability hypothesis, widespread species from resource rich habitats where more damaged by herbivores (less defended) than widespread species from resource poor habitats – such a growth-defense tradeoff was lacking in rare species. This indicates that the evolutionary important tradeoff between traits increasing competitive-ability and defence is present in widespread species but may have been lost in rare species.

**Main conclusions:** Our results indicate that biotic interactions, above all competition, might indeed set range limits, and underlines the importance of including context-dependency in studies comparing traits of common and rare or invasive and non-invasive species.

## Introduction

Understanding why some species are rare while others are widespread or invasive remains a fascinating question in ecology spanning decades (Baker, 1965; Gaston, 1994). To identify the factors which cause species to be more or less successful than others, in plants, studies comparing rare and widespread species, or invasive and non-invasive species, have a long tradition. While early attempts were often restricted to few species (Gaston, 1994; Murray, Thrall, Gill, & Nicotra, 2002), in recent times phylogenetically controlled multi-species experiments provided powerful tests for general patterns in rarity or invasiveness (Dawson, Fischer, & van Kleunen, 2012; Van Kleunen, Weber, & Fischer, 2010; van Kleunen, Dawson, Bossdorf, & Fischer, 2014). However, attempts to identify traits or characteristics of rare and widespread or invasive species were often not successful or revealed contradictory results. One reason for this is that certain plant traits or characteristics may lead to plant success in some habitats, but may be costly (as they often trade off, Kempel *et al.* 2011) or unimportant in others (Darwin 1859; Louthan *et al.* 2015). This context dependency was frequently invoked to explain that the drivers of plant success change along environmental gradients (Funk & Cornwell, 2013; Kueffer, Pyšek, & Richardson, 2013; van Kleunen, Dawson, & Maurel, 2015). However, context dependency has rarely been considered in experimental studies comparing characteristics of plants of widespread and rare, or invasive and non-invasive species.

Particularly biotic interactions, which can have strong effects on a species fundamental niche and may lead to plant rarity or invasiveness (Pigot & Tobias, 2013; van der Putten, Macel, & Visser, 2010; Wisz et al., 2013), are highly context dependent. However, they are rarely considered in studies explaining rarity and commonness (Brown, 1984; Hanski, Kouki, & Halkka, 1993). For example, the strength of competition between plants strongly depends on resource availability - in resource-rich habitats, above- and belowground competition are strong (Tilman, 1988), whereas in resource-poor habitats plants compete mainly for belowground resources, relaxing the strength of competitive exclusion (Grime, 1977; Hautier, Niklaus, & Hector, 2009; Tilman, 1988) or even promoting facilitation among plants (Callaway et al., 2002). Concordantly, competition might be important for a plant’s success in resource rich environments, whereas in resource limited environments success might be rather driven by abiotic factors (Connell, 1961; Louthan et al., 2015; Warren & Bradford, 2011). Many widespread species occupy nutrient-rich environments and often have adaptations for high resource acquisition, whereas many regionally rare species are characterized by resource-conservatism, and are thus limited to resource-poor environments (Drury, 1974; Grime, 1977). This has led to the long standing hypothesis that widespread species are competitively superior over rare ones (Griggs, 1940; Powell & Knight, 2009). However, if we compare the competitive ability of widespread and rare species, and by chance most rare species originate from resource poor and most widespread species from resource rich habitats, results may be biased and simply reflect differences in the importance of competition along a resource axis (Dawson et al., 2012; Lloyd, Lee, & Wilson, 2002). Therefore, it is crucial to take the position of a species along a resource gradient into account, and to compare rare and common species originating from both, resource poor and rich habitats, if we want to adequately test for differences in the competitive ability of widespread and rare species.

Similarly, the impact of plant enemies such as herbivores, and hence the importance of a plant to defend against them, or to tolerate them, may strongly depend on environmental context. Species which evolved in resource-poor environments are suggested to be less able to replace lost tissue and should thus invest more into defences than faster-growing, competitive species from more productive environments which can rapidly compensate biomass loss (growth-rate hypothesis, Coley, Bryant, & Chapin, 1985). In line with this, plants from productive environments should be more tolerant to herbivory or pathogen infestation than plants from resource poor ones. In addition, there are also good reasons to believe that regionally rare plant species in general are less defended against enemies than widespread species, although this has been rarely tested (but see Kempel et al. 2018). Rare species often occur in small or isolated populations, and may thus have a low genetic diversity, making them more susceptible to enemies (Spielman, Brook, Briscoe, & Frankham, 2004). Alternatively, regionally rare plant species are often less apparent to their enemies than more widespread ones, and fewer herbivores or pathogens might have specialized on them (Feeny, 1976). Such a reduced apparency to enemies could have led to the evolution of reduced defence, making rare plant species more susceptible to herbivory or pathogen attack if subsequently exposed (Laine, 2006). Consequently, if we compare rare and widespread plant species in their ability to resist or tolerate their enemies, and by chance most rare species originate from resource poor and most widespread plant species from resource rich habitats, our results may not reflect differences in rarity, but rather differences in resource deployment strategies along a resource gradient. Again, this highlights the importance of the inclusion of context dependency in studies comparing characteristics of rare and widespread plant species.

Here, we present a multispecies experiment where we compared the competitive ability, the tolerance to herbivory (i.e. the regrowth capacity to experimental clipping) and ambient herbivore damage (as an indication of plant resistance) between 19 regionally rare and endangered and 21 widespread plant species from Switzerland. To make sure that the differences we present are due to regional rarity and are not confounded with local plant rarity (abundance), we assured that the same amount of rare and widespread species usually reach high abundances at a local scale, and the same amount usually occur in low abundances locally (Table 1). The rare and widespread species were selected from the same genus or plant family to account for phylogenetic influences, and each species group originated from a similar habitat (Table 1). Habitats between species groups differed greatly in their resource availability. This allowed us to test for differences in the ability to cope with antagonistic biotic interactions of many widespread and rare species while accounting for variation in environmental and phylogenetic context. Additionally, because the outcome of competition and simulated herbivory may interact with each other, and might depend on the resource conditions of the experiment, we applied a nutrient addition treatment and assessed any relationship with plant rarity or a plants position along a resource gradient and all factors interactively.

**Table 1:**
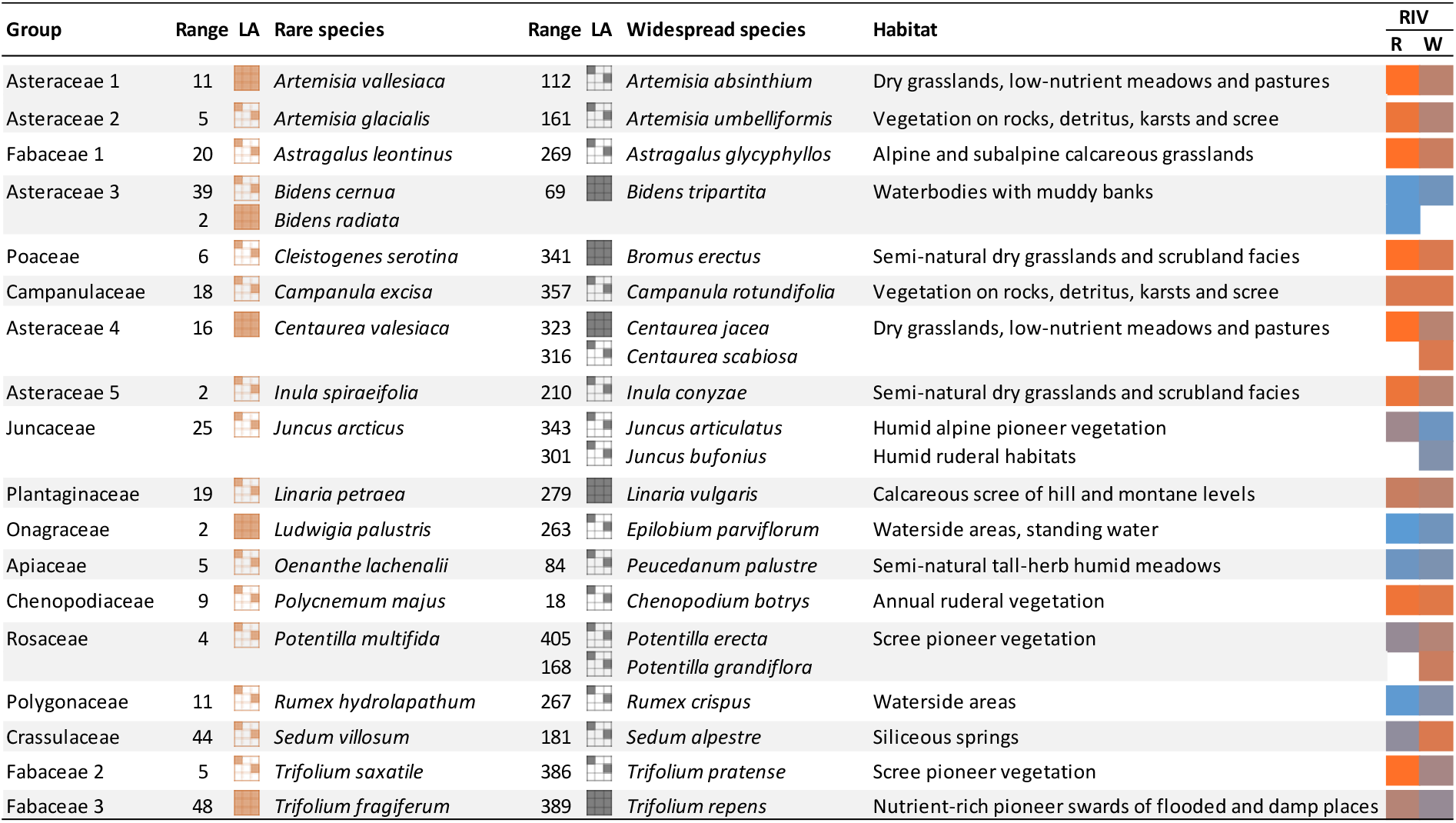
Species selected for the experiment. Species were grouped according to phylogeny and habitat such that each group consists of at least one regionally rare and one widespread species from the same genus or family and occupying the same or a similar habitat. The range size (Range) for each species is given as the number of 10 × 10 km grid cells occupied by a given species in Switzerland. The local abundance (LA) of each species is depicted with symbols and indicate whether species usually grow in larger groups or stands at the place where they occur (locally abundant, 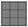), or whether they are usually scattered or only grow in small groups (locally scattered, 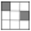). Resource indicator values (RIV) for rare (R) and widespread species (W) are depicted using a colour code with red and blue colours indicating dry and nutrient poor, respectively wet and nutrient rich conditions.

| Group | Range | LA | Rare species | Range | LA | Widespread species | Habitat | RIV |  |
| --- | --- | --- | --- | --- | --- | --- | --- | --- | --- |
|  |  |  |  |  |  |  |  | R | W |
| Asteraceae 1 | 11 |  | *Artemisia vallesiaca* | 112 |  | *Artemisia absinthium* | Dry grasslands, low-nutrient meadows and pastures |  |  |
| Asteraceae 2 | 5 |  | *Artemisia glacialis* | 161 |  | *Artemisia umbelliformis* | Vegetation on rocks, detritus, karsts and scree |  |  |
| Fabaceae 1 | 20 |  | *Astragalus leontinus* | 269 |  | *Astragalus glycyphyllos* | Alpine and subalpine calcareous grasslands |  |  |
| Asteraceae 3 | 39 |  | *Bidens cernua* | 69 |  | *Bidens tripartita* | Waterbodies with muddy banks |  |  |
|  | 2 |  | *Bidens radiata* |  |  |  |  |  |  |
| Poaceae | 6 |  | *Cleistogenes serotina* | 341 |  | *Bromus erectus* | Semi-natural dry grasslands and scrubland facies |  |  |
| Campanulaceae | 18 |  | *Campanula excisa* | 357 |  | *Campanula rotundifolia* | Vegetation on rocks, detritus, karsts and scree |  |  |
| Asteraceae 4 | 16 |  | *Centaurea valesiaca* | 323 |  | *Centaurea jacea* | Dry grasslands, low-nutrient meadows and pastures |  |  |
|  |  |  |  | 316 |  | *Centaurea scabiosa* |  |  |  |
| Asteraceae 5 | 2 |  | *Inula spiraeifolia* | 210 |  | *Inula conyzae* | Semi-natural dry grasslands and scrubland facies |  |  |
| Juncaceae | 25 |  | *Juncus arcticus* | 343 |  | *Juncus articulatus* | Humid alpine pioneer vegetation |  |  |
|  |  |  |  | 301 |  | *Juncus bufonius* | Humid ruderal habitats |  |  |
| Plantaginaceae | 19 |  | *Linaria petraea* | 279 |  | *Linaria vulgaris* | Calcareous scree of hill and montane levels |  |  |
| Onagraceae | 2 |  | *Ludwigia palustris* | 263 |  | *Epilobium parviflorum* | Waterside areas, standing water |  |  |
| Apiaceae | 5 |  | *Oenanthe lachenalii* | 84 |  | *Peucedanum palustre* | Semi-natural tall-herb humid meadows |  |  |
| Chenopodiaceae | 9 |  | *Polycnemum majus* | 18 |  | *Chenopodium botrys* | Annual ruderal vegetation |  |  |
| Rosaceae | 4 |  | *Potentilla multifida* | 405 |  | *Potentilla erecta* | Scree pioneer vegetation |  |  |
|  |  |  |  | 168 |  | *Potentilla grandiflora* |  |  |  |
| Polygonaceae | 11 |  | *Rumex hydrolapathum* | 267 |  | *Rumex crispus* | Waterside areas |  |  |
| Crassulaceae | 44 |  | *Sedum villosum* | 181 |  | *Sedum alpestre* | Siliceous springs |  |  |
| Fabaceae 2 | 5 |  | *Trifolium saxatile* | 386 |  | *Trifolium pratense* | Scree pioneer vegetation |  |  |
| Fabaceae 3 | 48 |  | *Trifolium fragiferum* | 389 |  | *Trifolium repens* | Nutrient-rich pioneer swards of flooded and damp places |  |  |

## Methods

### Plant species

To test whether rare and widespread plant species respond differently to biotic interactions, we originally selected 52 plant species from a wide range of different habitats. These species differ on the one hand in their degree of regional rarity and endangerment in Switzerland and on the other hand, in the resource availability of the habitat they typically occupy. We classified 27 plant species as ‘rare’ and 25 plant species as ‘widespread’ *a priori*. Rare species were listed as near-threatened, vulnerable, endangered or critically endangered according to the Red list for Switzerland (Moser, Gygax, Bäumler, Wyler, & Palese, 2002). Widespread species were listed as being of least concern except for three species, which were listed as near-threatened, but which had a relatively large range size (and particularly larger than their rare congeneric partner, Table 1). We calculated the range size of these species as the number of 10 × 10 km grid cells occupied in Switzerland and found that rare species had 94% smaller range sizes than widespread species (F_1,38_ = 82; P < 0.0001, analysis done on the final selection of species). Next, to control for phylogenetic and habitat effects, we grouped rare and widespread species, so that groups contained at least one rare and one widespread species from the same genus or plant family and from the same habitat, resulting in 21 groups. Five species did not germinate in sufficient numbers and were excluded together with their congeneric partner from the analysis, ending up with 21 common and 19 rare species, forming 18 groups (Table 1). To ensure that the differences we report here are due to regional rarity and not local plant abundance, for both the rare and the widespread species, we selected some species that usually reach high abundances at a local scale, and some species that usually occur in low abundances at the place where they occur (“dominance in situ” from the Flora Indicativa, Landolt et al.(2010), values of our species range from 1-4, with 1 = scattered, 2 = scattered or in small groups, 3 = in larger groups, 4 = in larger stands, Table 1).

The habitats of our selected species differed mainly in their soil moisture and nutrient availability. Thus, to describe the position of the realized niche optima of our species along a resource gradient, we used the ecological indicator values for nutrients and moisture after Landolt et al. (Flora Indicativa, 2010). Landolt indicator values follow an ordinal scale and range from 1-5 (low numbers represent low, high numbers high nutrient and moisture requirements) and have been widely used in ecological studies. Nutrient and moisture values were correlated (r = 0.33, P = 0.017). To avoid collinearity but maintain full environmental space, we calculated a principal component analysis, and used the scores of the first axis for further analysis, hereafter named resource indicator value (explained variance = 72%, values ranged from −1.89 to 2.19 with high values indicating that species originate from wet and nutrient rich habitats, Fig. S1, Table 1). Rare and widespread plant species did not differ in their resource indicator value (F_1,38_ = 0.504, P = 0.48). Seeds of rare plant species were collected in the wild (seeds of one population per species of at least 10 mother plants), seeds of widespread plant species were either collected in the wild (seeds of one population per species of at least 10 mother plants) or obtained from commercial seed suppliers.

### Experimental design

In spring 2013 we sowed seeds of our species in trays. After germination we transplanted 40 seedlings per species individually into 1.3 l pots filled with mixed soil containing 20% compost, 20% agricultural field soil from Swiss Plateau region, 20% wood fibre, and 40% peat. Pots were then placed outside in a common garden at Muri bei Bern (46.9351° N, 7.4985° E, Switzerland). To test whether the species respond differently to biotic interactions at different levels of resources, we applied a fully factorial experiment, including a competition treatment, a clipping treatment to simulate herbivory and to test for regrowth capacity, and a fertilizer treatment to change resource supply experimentally. This allowed us to test how the response to competition and simulated herbivory changes with increasing resource supply and whether these responses differed between widespread and rare species. We applied all possible combinations of these treatments (40 species × 8 treatments × 5 replicates = 1600 plants), however, for some species we did not always have 5 replicates per treatment or could not perform all treatment combinations (1351 plants in total, Supporting Table S1). Pots were then distributed among 5 blocks (one replicate per species and treatment per block) and pots receiving the same treatments were grouped together to facilitate the applications of the treatments.

To simulate competition, we sowed 1g of a common grass species, *Lolium perenne*, in pots of the competition treatment 5 weeks after transplanting seedlings. Competition with *L. perenne* may not reflect competition against all natural competitors of our rare and widespread species, however, we wanted to have the same competitor for all species to be able to compare the results, and choose *L. perenne* as it is likely to not co-occur with any of our species. At the same time, to simulate herbivory we removed ca. 50% of the leaf biomass of the target plants by clipping each leaf by half for herbs, or in the middle for grasses. To experimentally increase resource supply, we fertilized plants every 2 weeks with a soluble NPK fertilizer (Wuxal).

To get an indication of a plant’s herbivore resistance we visually recorded damage caused by ambient leaf chewing herbivores in the common garden on all plant species (percentage herbivore damage = number of infested leaves × percentage damage of infested leaves / total number of leaves). In August 2013 we harvested aboveground biomass of all target plants, and of the competitor **Lolium perenne**. Biomass was dried (72h at 70°C) and weighed.

### Statistical analysis

We used linear mixed effects models (lmer, package lme4 in R; R Core Team 2013). To assess whether regionally rare plant species suffered more from competition or experimental clipping than widespread plant species, and whether this changes with fertilization we used aboveground biomass (log-tranformed) of our target species as response variable and fitted status (widespread, rare), competition, clipping, fertilization and all possible combinations as fixed terms. To test whether the response to the treatments and differences between widespread and rare species depended on the resource availability of their habitat, we additionally fitted the species resource indicator value and all interactions with all other factors as fixed effects (see Table S3). Grasses or annuals may respond differently to competition or clipping than herbaceous or perennial plants. To account for this, we included functional group (grass, herb) and lifeform (not perennial, perennial) in our model and all possible interactions, except higher-order interactions that involve status, resource indicator value, functional group or lifeform which we had to remove, because we had too few grasses and annuals in our study to test for these interactions.

We also tested whether the effect of our plants on the biomass of the competitor *L. perenne* depended on rarity, resource origin, clipping or fertilizer, and used aboveground biomass (log-transformed) of *L. perenne* as response variable and all factors and interactions as above (except those involving competition) as fixed terms.

Further, we tested whether the amount of ambient herbivory was related to rarity, resource indicator value and all treatments, and used the percentage leaf damage (arcsin square root transformed) as response variable and included the same factors as in the model for aboveground biomass as fixed terms.

In the model for the biomass of the competitor *L. perenne* we either included or excluded the final biomass of the target species as a covariable. In all models we included species (40 levels) nested into groups (18 levels, congeneric or confamilar, Table 1) and plant family (12 levels), and block (five levels) as random terms. We simplified the best model (but kept the main factors status, resource indicator value, clipping, competition and fertilization in the models) and derived significances using likelihood-ratio tests comparing models with and without the factor of interest.

## Results

Overall, rare plant species had a lower biomass than widespread plant species (Fig. 2, Status: Chi^2^ = 4.07, *P*=0.044, Table S2). Competition with *L. perenne* and simulated herbivory by experimental clipping reduced plant biomass (Competition: Chi^2^=10.4, P=0.001; clipping: Chi^2^=82.22, P < 0.0001), whereas fertilization alone had no effect on plant biomass (Table S2). In general, common and rare plant species did not differ in their response to competition (no significant Competition × Status interaction), their response to experimental clipping (no significant Clipping × Status interaction) nor in their response to fertilization (no significant Fertilizer × Status interaction, Table S2).

Grasses and herbs and annual and perennial species did not differ in their aboveground biomass (Functional Group and Lifeform not significant, Table S2). However, annual plants were less affected by competition than perennial plants (Competition × Lifeform interaction: Chi^2^= 5.25, P=0.022, Table S2).

The effect of competition depended on the resource indicator value (Competition × Resource origin interaction: Chi^2^= 14.56, P=0.0001, Table S2), whereas the effect of clipping and fertilization did not depend on the resource indicator value. Competition had especially strong negative effects on species with a high resource indicator value. When species had a high resource indicator value, i.e. originated from nutrient rich and moist habitats, rare plant species suffered more from competition than common plant species (Fig. 1, Status × Competition × Resource indicator interaction: Chi^2^= 9.03, P= 0.003, Table S2). In contrast, for plants with a low resource indicator value, i.e. plants originating from nutrient poor and dry habitats, common and rare species did not differ in their response to competition.

**Figure 1:**
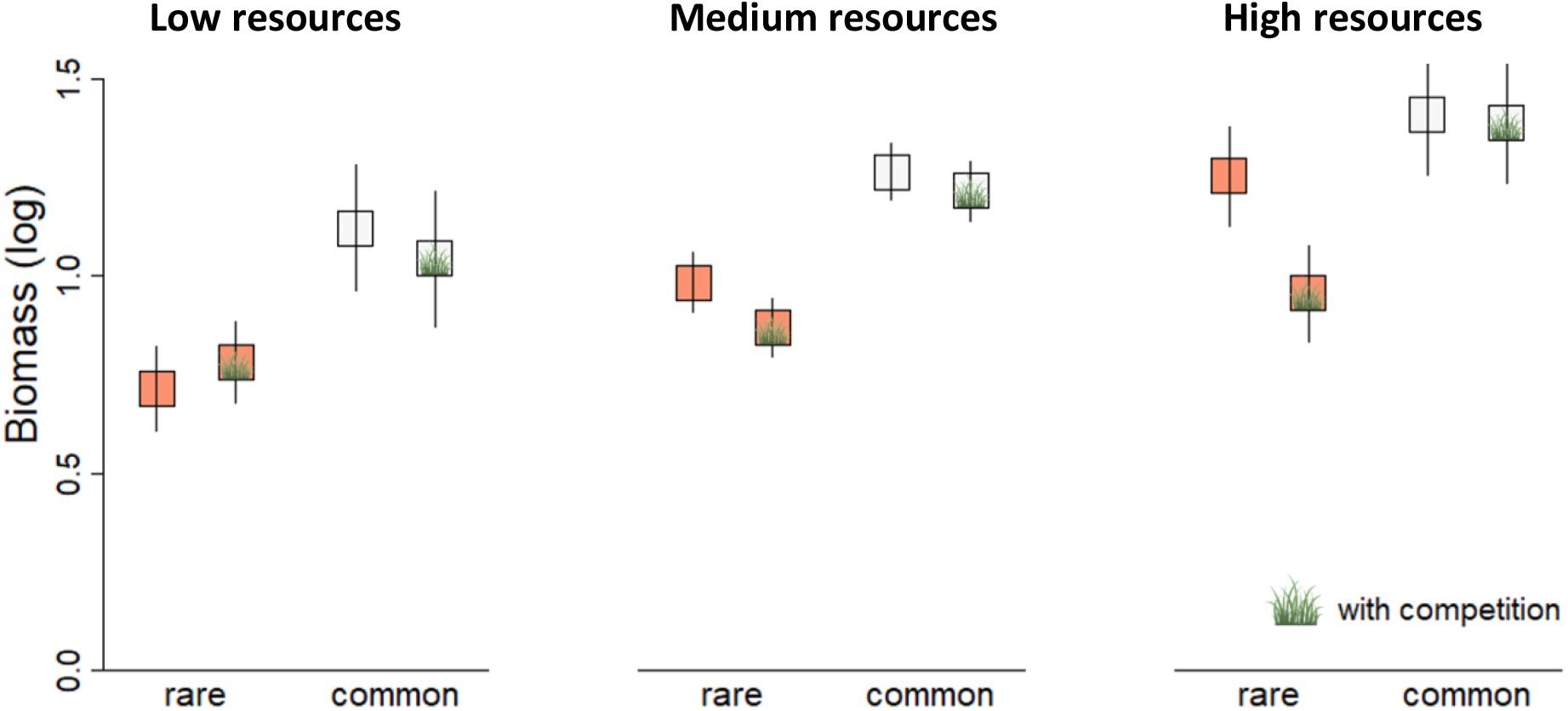
Aboveground biomass (log-transformed, in g) of rare and regionally common plant species without and with competition originating from low, medium or high resource habitats (resource indicator values −1.7, 0.15 and 2 respectively). Shown are fitted estimates from a liner mixed effect model. Error bars indicate confidence intervals (obtained from the effect package in r).

**Figure 2:**
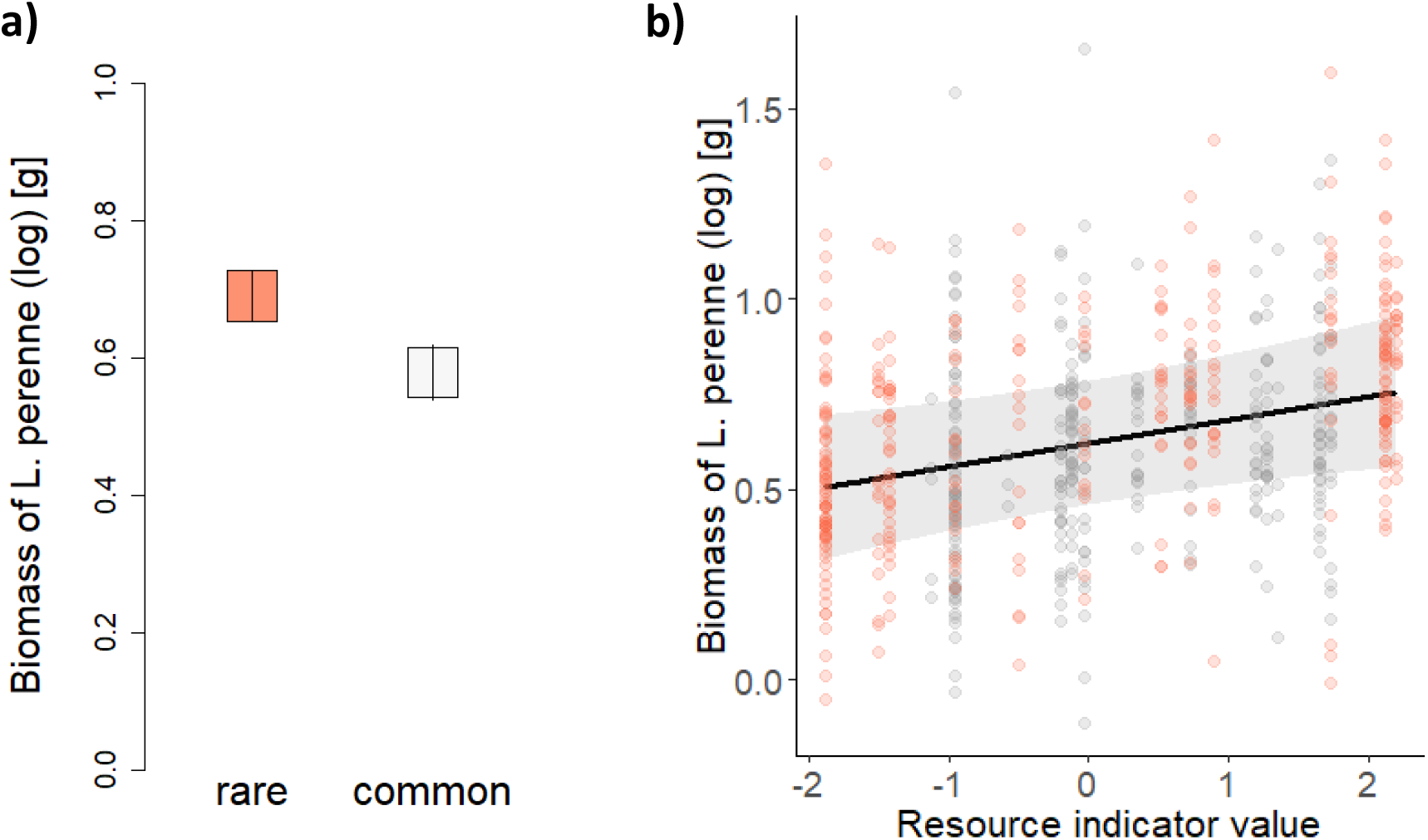
Biomass of the competitor *L. perenne* growing with a) rare and regionally common plant species, when the biomass of the target species was not included as a covariable, and with b) both rare and common plant species originating from habitats of different nutrient and moisture values, when their biomass was included as a covariate in the model. High values of resource indicator values indicate that a species originate from nutrient rich and moist habitats, low values indicate that a species originate from nutrient poor and dry habitats. Shown are the fitted values, respectively lines, and partial residuals from a linear mixed effect model. The error bars, respectively shaded area, indicate lower and upper confidence intervals (obtained from the effect package in r). The points are partial residuals obtained from the visreg package (red=rare species, grey=regionally common species). When we accounted for the biomass of the target species, biomass of *L.perenne* did not significantly differ between regionally common and rare species.

When plants had been fertilized the combined effects of competition and clipping had the strongest negative effects on plant biomass, whereas in non-fertilized pots clipping and competition combined had as negative effects on plant biomass as clipping alone (Fertilizer × Competition × Clipping interaction: Chi^2^= 4.62, P = 0.032, Table S2 and Fig. S2), indicating that biomass loss due to herbivory or mowing can be tolerated easier under high nutrient conditions, where resources for a fast regrowth are more abundant.

Further, fertilization alleviated the negative effects of clipping, but only for plants with a high resource indicator value (Fertilization × Clipping × Resource indicator interaction: Chi^2^=5.17, P=0.023). Clipping similarly reduced biomass of unfertilized and fertilized plants with a low resource indicator value, whereas for plants with a high resource indicator value, the negative effect of clipping was smaller when plants had been fertilized (Fig. S3). This indicates that plants from resource rich origins are the ones that can better tolerate biomass loss under high nutrient conditions, potentially due to their better ability to monopolize belowground resources for regrowth. However, this pattern might also arise because our unclipped plants could not profit from fertilization, potentially because they were already pot-bound. We would thus interpret the finding that species from resource rich habitats are better able to tolerate herbivory under high nutrient conditions with caution.

The biomass of the competitor *Lolium perenne* was higher when it was growing with rare plant species compared to common plant species, indicating that common species had a stronger competitive effect on *L. perenne* (Status: Chi^2^= 4.95, P = 0.026, Table S3, Fig 2). When we included biomass of the target species as a covariable, common and rare species only marginally differed in their effect on *L. perenne*, indicating that the stronger negative effect of common species is mainly driven by their higher aboveground biomass (Status: Chi^2^= 3.16, P= 0.075, Table S3). In both models, the biomass of *L. perenne* was lower when neighbouring plants had a low resource indicator value (i.e. originated from nutrient poor and dry habitats) than when they had a high resource indicator value (Resource origin: Chi^2^= 4.11, P=0.042) and this effect became stronger when we accounted for differences in aboveground biomass of the target species (Resource indicator value, with biomass of the target plants as covariable: Chi^2^= 7.23, P=0.007, Table S3, Fig. 2). This might indicate that for *L. perenne*, belowground competition with species from resource poor habitats is stronger than with species from the resource rich end.

Overall, percentage of herbivore damage was not affected by competition and fertilization, and was reduced when plants were clipped (Table S4). Herbivore damage did not differ between common and rare plant species (Status: Chi^2^= 0.20, P=0.65, Table S4), and was not related to the resource indicator value of a species (Resource origin: Chi^2^= 0.051, P=0.82). However, the percentage of herbivore damage increased with a species’ resource indicator value in common species, whereas in rare species the percentage of herbivore damage did not change with a species’ resource indicator value (Chi^2^= 4.93, P=0.026, Fig. 3). This indicates a tradeoff between fast growth and being competitive on one hand and defence on the other hand in common species but no such tradeoff in rare species, which showed intermediate levels of herbivore damage, independent of their position along a resource gradient.

**Figure 3:**
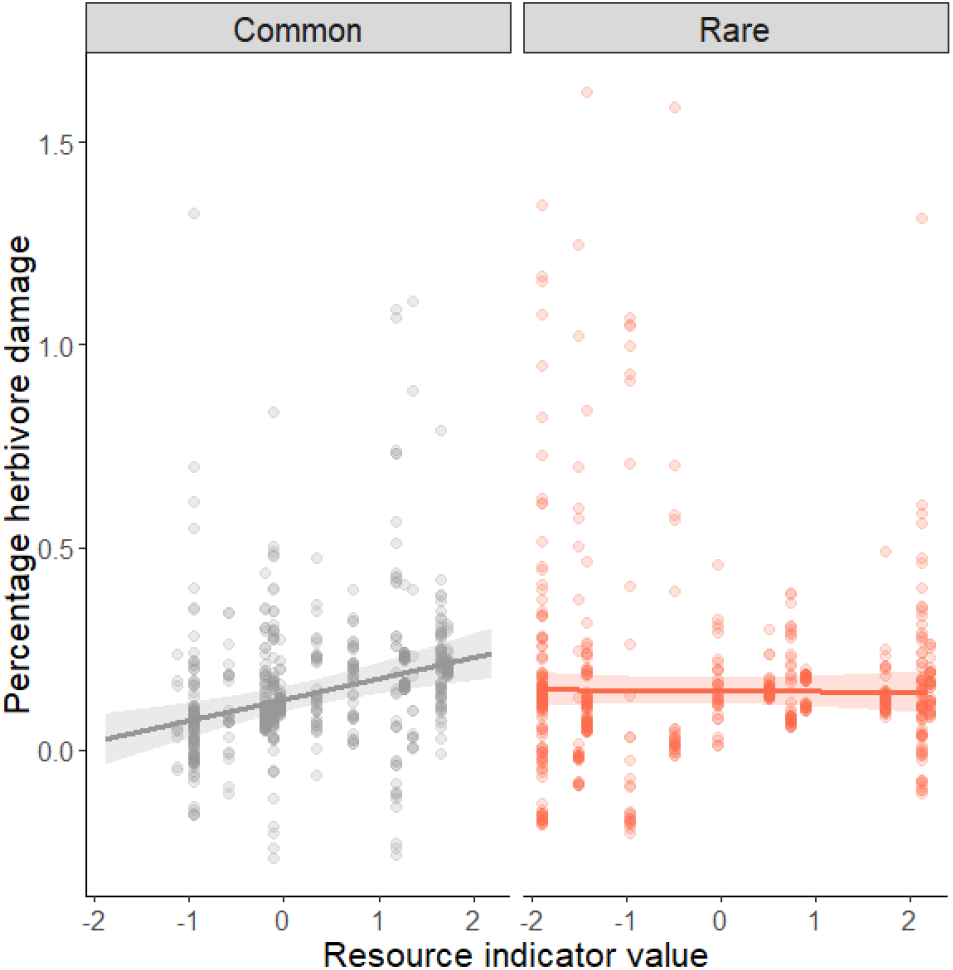
Percentage herbivore damage (arc-sinus square root transformed) on rare and regionally common plant species with different resource indicator values. High resource indicator values indicate that a species originate from nutrient rich and moist habitats, low values indicate that a species originate from nutrient poor and dry habitats. Shown are the fitted lines from a liner mixed effect model and the standard errors (obtained from the effect package). Points are partial residuals obtained from the visreg package.

## Discussion

Although biotic interactions are increasingly recognized as important factors in determining species range limits (Pigot & Tobias, 2013; van der Putten et al., 2010; Wisz et al., 2013), evidence that rare and widespread species respond differently particularly to antagonistic biotic interactions is still scarce and often controversial, potentially because the importance of certain plant characteristics for a plant’ success is context dependent. In our multi-species experiment, using 40 plant species differing in regional rarity and originating from contrasting habitats, we show that taking into account phylogenetic and environmental context dependency is crucial when comparing widespread and rare or invasive and non-invasive species.

## Competitive ability, rarity and resource origin

Although the importance of interspecific competition along a resource gradient has long been debated (Grace, 1991; Grime, 1979; Tilman, 1988), it seems obvious that plant characteristics leading to success in resource rich habitats differ from those leading to success in resource limited habitats. This is because resources for which species are competing differ, mainly light in nutrient rich and moist habitats and nutrients and water in nutrient limited and dry habitats (Wilson & Tilman 1991, Connell 1961; Grime 1979). Accordingly, characteristics of species from resource rich habitats comprise traits leading to fast growth and rapid capture of light and belowground resources whereas species from resource-limited habitats possess traits leading to nutrient retention. Both adaptive strategies are suggested to trade off (Aerts, 1999; Reich, 2014).

Having this is mind it is not surprising that studies comparing the competitive ability of rare and common species or invasive and non-invasive alien species without considering their ecological context often find controversial results (e.g. Rabinowitz et al. 1984; Powell & Knight 2009; Dawson et al. 2012). For instance, in an experiment comparing the competitive response of native rare and common and alien rare and common species, Dawson et al. (2012) found a tendency for higher survival under competition in more common (native or alien) species, but this effect disappeared when they corrected for the fact that most rare species originated from resource poor and most common species from resource rich habitats.

In our experiment, using 40 plant species differing in regional rarity and originating from a wide range of habitats, we show that widspread species suffered less from competition than rare species, but only when both widespread and rare species originated from nutrient rich and moist habitats. Instead, when species originated from more nutrient limited and drier habitats, widespread and rare species were similarly affected by competition. At least for species originating from habitats where aboveground competition for light is usually strong our results support the long-standing hypothesis that common species are competitively superior compared to rare species (Griggs, 1940). It is thus likely that a higher competitive ability in these habitats have helped them to expand their ranges and to become common, and that competition could potentially set range limits in abiotically less stressful conditions.

For species from resource poor and dry habitats, a high ability to tolerate low levels of nutrients and water rather than a high competitive ability for light is important. In our experiment, fertilization surprisingly did not have a consistent positive effect, indicating that nutrients were not limited. Thus, we cannot test whether widespread species from resource-limited habitats would have performed better under dry and nutrient limited conditions than rarer species. Whether certain species from resource poor environments are widespread because they can better tolerate low levels of nutrients and dry conditions, or whether they are stronger competitors for belowground resources, remains to be resolved. For species from the resource rich end, however, it seems that indeed the outcome of biotic interactions such as competition could be an important driver of large-scale plant rarity and is likely to determine range limits of species.

Not only the response to competition but also the ability of a plant to supress the growth of other plants – the competitive effect – is a component of the competitive ability of species (Goldberg, 1990). Widespread species in our experiment generally had stronger negative effects on the competitor *L. perenne* than rare species. This was mainly because widespread species had a higher aboveground biomass, which is consistent with previous studies (Cornwell & Ackerly, 2010; Dawson et al., 2012; Lavergne, Garnier, & Debussche, 2003; Murray et al., 2002). Interestingly, whether a plant had a strong or weak effect on the growth of *L. perenne* also depended on whether it originated from a resource poor or rich habitat: plants from nutrient poor and dry habitats had stronger negative effects on *L. perenne* than plants originating from nutrient rich and moist habitats. Likely, this is caused by traits related to stronger belowground competition such as a larger root system of plants from resource poor compared resource rich environments (Tilman 1988), but also different allelopathic activities or different microbial communities around roots of species from contrasting environments could potentially drive this pattern (Inderjit, Wardle, Karban, & Callaway, 2011; Kempel, Rindisbacher, Fischer, & Allan, 2018). In summary, widespread species from resource rich environments not only suffer less from competition than rare species, but in general widespread species have stronger negative effects on other plants due to their larger biomass, which might both have contributed to their large ranges.

## Tolerance and resistance to herbivory, plant rarity and resource origin

Herbivores have been shown to have strong effects on the relative abundance and composition of species in communities (e.g. McNaughton 1979; Allan & Crawley 2011), however whether and how they can determine patterns of large scale plant rarity is questionable. Plants can cope with herbivores by increasing their tolerance or their resistance (Karban & Baldwin, 1997), and although variation in these attributes differ greatly between species, they have rarely been related to large-scale rarity.

We did not find that widespread and rare species respond differently to experimentally simulated herbivory, and thus that tolerance is related to large scale patterns of plant rarity. Also, whether a species originated from resource poor or rich habitats did not generally affect its regrowth capacity, although there was an interaction of fertilization, clipping and resource indicator value. This showed that fertilization could alleviate the negative effects of clipping, but only for those plants that originated from nutrient rich and moist habitats (Fig. S3). However, this pattern might have arisen because our fertilizer treatment did not show the expected positive effect on unclipped plants, likely because these plants were already pot-bound due to their high biomass.

How abiotic or biotic factors affect tolerance to herbivory is still largely unclear (Fornoni, 2011; Strauss & Agrawal, 1999; Wise & Abrahamson, 2007). In accordance to the resource availability hypothesis (Coley et al., 1985) fast growing species with rapid rates of nutrient absorption from nutrient rich environments have been found to have high levels of tolerance (Gianoli & Salgado-Luarte, 2017). Others found that nutrient availability is negatively related to tolerance (see Strauss & Agrawal 1999, Wise & Abrahamson 2007), because high nutrient levels reduce the amount of roots relative to shoots, which in turn reduces regrowth capacity. Species with effective nutrient retention and a large root system, which are often found in resource poor environments, should thus be more tolerant. Since we found no relationship between plant tolerance and the position of our species along a resource axis, it is likely that particularly under nutrient rich conditions as we had in our experiment, both strategies – rapid rates of nutrient uptake and a large root system - led to high levels of plant tolerance. In conclusion, our study indicates that plant tolerance alone does not seem to be related to large-scale plant rarity and commonness when species habitat characteristics are taken into account, likely because tolerance together with resistance are both strategies for plants to cope with herbivory.

Only few studies tested whether widespread or rare plants differ in their resistance to herbivores or other plant enemies. Landa & Rabinowitz (1983) found that a common grasshopper preferred rare species over more apparent and widespread grass species (7 species), Fiedler (1987) found that rare species are more prone to leaf grazing (4 species), supporting the idea that apparent plants are more defended. Similarly, Kempel et al. (2018) showed that rare plant species were more susceptible to soil biota than widespread species (19 species), indicating that defences are lower in rarer species.

In our study, we found no evidence that rare species are more susceptible to ambient leaf herbivory, and thus that they are less defended than widespread species. Interestingly, however, in widespread plants the level of herbivore damage depended strongly on whether plants originated from resource rich or low habitats: consistently with the resource availability hypothesis plants from resource rich habitats were more damaged (indicating that they are less defended) than species from resource poor habitats. In contrast, rare species showed intermediate levels of damage over all resource origins. It thus seems as if the tradeoff between traits that increase competitive ability and defence is only present for widespread species – likely a successful adaptation to changes in the relative importance of herbivory and competition for plant success along a resource axis. In resource rich habitats a low investment in defences might allow plants to be competitively superior, particularly if plants are less limited by herbivores than by competing neighbouring plants. Rare species, which often have undergone bottlenecks and show low levels of genetic diversity (Gaston, 1994), might have lost such adaptations and this might have broken up tradeoffs (Kempel et al., 2011). Although this is only speculative, the potentially higher investment in defence for rare species from resource rich habitats might be one explanation for why they are less competitive compared to widespread species. So far, our data do not indicate that rare and widespread species generally differ in their tolerance and their susceptibility to herbivores. Nevertheless, it might be interesting to further investigate whether important evolutionary tradeoffs are lost in rare species.

## Conclusion

Our multi-species experiment controlling for phylogenetic and environmental context supports the long-standing hypothesis that regionally common species are better competitors than rare species, but restricts it to only those species originating from resource rich habitats, in which a high competitive ability is advantageous. Using many species from a large number of environmental conditions suggests that this is a general pattern. It is thus likely that competitive interactions might be important drivers of large-scale plant rarity. Our results clearly underline the importance of context dependency - much controversy in studies comparing characteristics of common and rare or invasive and non-invasive species could potentially be explained if we would consider that the importance of certain species characteristics for plant success might differ depending on environmental conditions.

Altogether, our data hint to the fact that biotic interactions, above all plant competition, are important drivers of species range limits. Increasing our understanding not only in environmental but also biotic drivers underlying species range distributions is therefore of utmost importance if we are to understand and predict species responses to global change, and to conserve todays biodiversity.

## Author’s contribution

AK, HV and MF designed the experiment, AK and HV performed research, AK analysed the data and wrote the manuscript with substantial input from DP, and all authors contributed substantially to revisions.

## Acknowledgements

We thank Karl Kaspar and Judith Hinderling for help with the common garden experiment, and the Federal Office for the Environment (FOEN) for funding.

## Data availability statement

Data will be archived in a public repository upon acceptance.

