## Supporting information for "Context dependency of biotic interactions and its relation to plant rarity"

**shared first-authorship*

**Correspondent author:** Anne Kempel, Institute of Plant Sciences, Altenbergrain 21, 3013 Bern, Switzerland,, +41 31631 4939

### Supporting Tables

Table S1: Number of plants per species and treatment. C = plants growing in competition with *L. perenne*, H = 50 % leaf biomass clipped to simulate herbivory, F = plants fertilized.

| Group | Speciesname | Control | C | H | F | CH | CF | FH | CFH |
|---|---|---|---|---|---|---|---|---|---|
| Asteraceae 1 | *Artemisia absinthium* | 5 | 5 | 5 | 5 | 5 | 4 | 5 | 5 |
| Asteraceae 2 | *Artemisia glacialis* | 5 | 4 | 5 | 4 | 4 | 5 | 5 | 3 |
| Asteraceae 2 | *Artemisia umbelliformis* | 4 | 3 | 3 | 4 | 4 | 4 | 2 | 0 |
| Asteraceae 1 | *Artemisia vallesiaca* | 5 | 5 | 5 | 5 | 5 | 4 | 5 | 5 |
| Fabaceae 1 | *Astragalus glycyphyllos* | 5 | 3 | 5 | 0 | 0 | 0 | 0 | 0 |
| Fabaceae 1 | *Astragalus leontinus* | 5 | 4 | 4 | 3 | 5 | 5 | 4 | 5 |
| Asteraceae 3 | *Bidens cernua* | 5 | 5 | 4 | 5 | 5 | 5 | 5 | 5 |
| Asteraceae 3 | *Bidens radiata* | 5 | 5 | 4 | 4 | 5 | 5 | 5 | 5 |
| Asteraceae 3 | *Bidens tripartita* | 5 | 5 | 4 | 4 | 5 | 5 | 4 | 5 |
| Poaceae | *Bromus erectus* | 4 | 5 | 5 | 5 | 3 | 4 | 5 | 2 |
| Campanulaceae | *Campanula excisa* | 4 | 5 | 5 | 5 | 4 | 5 | 5 | 5 |
| Campanulaceae | *Campanula rotundifolia* | 5 | 5 | 5 | 5 | 5 | 5 | 4 | 5 |
| Asteraceae 4 | *Centaurea jacea* | 4 | 3 | 4 | 4 | 5 | 5 | 4 | 5 |
| Asteraceae 4 | *Centaurea scabiosa* | 5 | 5 | 5 | 5 | 5 | 5 | 5 | 5 |
| Asteraceae 4 | *Centaurea valesiaca* | 4 | 5 | 5 | 5 | 5 | 5 | 3 | 5 |
| Chenopodiaceae | *Chenopodium botrys* | 5 | 5 | 5 | 0 | 0 | 0 | 0 | 0 |
| Poaceae | *Cleistogenes serotina* | 5 | 5 | 5 | 4 | 5 | 4 | 4 | 5 |
| Onagraceae | *Epilobium parviflorum* | 5 | 5 | 5 | 5 | 5 | 5 | 5 | 3 |
| Asteraceae 5 | *Inula conyzae* | 5 | 5 | 5 | 5 | 5 | 4 | 4 | 5 |
| Asteraceae 5 | *Inula spinosa* | 5 | 5 | 5 | 4 | 5 | 4 | 3 | 5 |
| Juncaceae | *Juncus arcticus* | 3 | 5 | 4 | 5 | 5 | 5 | 5 | 5 |
| Juncaceae | *Juncus articulatus* | 4 | 5 | 5 | 5 | 5 | 5 | 5 | 5 |
| Juncaceae | *Juncus bufonius* | 5 | 5 | 5 | 5 | 5 | 5 | 5 | 5 |
| Plantaginaceae | *Linaria alpina subs. petrea* | 5 | 5 | 4 | 4 | 4 | 5 | 5 | 5 |
| Plantaginaceae | *Linaria vulgaris* | 5 | 5 | 5 | 5 | 5 | 5 | 5 | 0 |
| Onagraceae | *Ludwigia palustris* | 4 | 5 | 5 | 5 | 5 | 5 | 5 | 5 |
| Apiaceae | *Oenanthe lachenalii* | 5 | 5 | 5 | 4 | 5 | 5 | 5 | 5 |
| Apiaceae | *Peucedanum palustre* | 5 | 5 | 0 | 5 | 0 | 0 | 0 | 0 |
| Chenopodiaceae | *Polycnemum majus* | 3 | 5 | 3 | 4 | 5 | 4 | 4 | 5 |
| Rosaceae | *Potentilla erecta* | 5 | 5 | 5 | 5 | 4 | 5 | 5 | 0 |
| Rosaceae | *Potentilla grandiflora* | 5 | 0 | 5 | 0 | 0 | 0 | 0 | 0 |
| Rosaceae | *Potentilla multifida* | 5 | 5 | 5 | 5 | 5 | 5 | 5 | 5 |
| Polygonaceae | *Rumex crispus* | 5 | 5 | 5 | 4 | 5 | 5 | 4 | 5 |
| Polygonaceae | *Rumex hydrolapathum* | 4 | 5 | 5 | 5 | 5 | 5 | 5 | 5 |
| Crassulaceae | *Sedum alpestre* | 4 | 5 | 5 | 5 | 5 | 5 | 4 | 5 |
| Crassulaceae | *Sedum villosum* | 4 | 5 | 5 | 1 | 5 | 5 | 5 | 5 |
| Fabaceae 3 | *Trifolium fragiferum* | 5 | 5 | 5 | 4 | 5 | 5 | 5 | 0 |
| Fabaceae 2 | *Trifolium pratense* | 5 | 5 | 5 | 4 | 5 | 4 | 5 | 4 |
| Fabaceae 3 | *Trifolium repens* | 5 | 5 | 5 | 5 | 4 | 5 | 5 | 5 |
| Fabaceae 2 | *Trifolium saxatile* | 5 | 4 | 0 | 0 | 0 | 0 | 0 | 0 |

Table S2: Results of a linear mixed effect model testing for the effects of biotic stresses and fertilization, rarity status and resource indicator value on plant biomass. Significances were obtained by stepwise deletion of non-significant terms and comparing the resulting model to the previous ones using log-likelihood-ratio tests. This resulted in a minimal model containing only significant terms (black). We kept random factors in the model and present their variances. Number in bold indicate statistical significance. To obtain $Ch^2$ and *P*-values of two-way interactions and main effects, we excluded all three-way, respectively all higher-order interactions and compared this model with models missing the factors of interest (indicated by [1)], respectively [2)]).

| Fixed factors | AIC | *Chi 2* | *P* | |
|---|---|---|---|---|
| Status | 1669.1 | 4.07 | 0.044 | 2) |
| Fertilizer | 1668.8 | 3.10 | 0.078 | 2) |
| Clipping | 1742.3 | 76.60 | <0.0001 | 2) |
| Competition | 1676.2 | 10.50 | 0.001 | 2) |
| Resource indicator value (RI) | 16668.2 | 2.45 | 0.117 | 2) |
| Functional group (FG) | 1614.58 | 0.05 | 0.727 | |
| Lifeform (LF) | 1666.5 | 0.79 | 0.373 | 2) |
| Status x Resource indicator value | 1625.4 | 0.01 | 0.913 | 1) |
| Status x Fertilizer | 1632.93 | 0.86 | 0.377 | |
| Status x Competition | 1627.6 | 2.25 | 0.133 | 1) |
| Status x Clipping | 1620.97 | 0.02 | 0.911 | |
| Status x Lifeform | 1667.84 | 0.52 | 0.471 | |
| Status x Functional group | 1656.15 | 0.39 | 0.532 | |
| Fertilizer x Resource indicator value | 1625.4 | 0.00 | 0.979 | 1) |
| Fertilizer x Lifeform | 1626.35 | 0.58 | 0.281 | |
| Fertilizer x Functional group | 1641.9 | 0.84 | 0.288 | |
| Fertilizer x Competition | 1625.6 | 0.25 | 0.617 | 1) |
| Fertilizer x Clipping | 1630.6 | 5.20 | 0.023 | 1) |
| Competition x Resource indicator value | 1640 | 14.56 | 0.0001 | 1) |
| **Competition x Lifeform** | **1651** | **5.10** | **0.023** | |
| Competition x Functional group | 1615.47 | 0.58 | 0.449 | |
| Clipping x Resource indicator value | 1626.4 | 1.03 | 0.311 | 1) |
| **Clipping x Lifeform** | **1651.7** | **4.94** | **0.026** | |
| Clipping x Functional group | 1619.01 | 0.03 | 0.738 | |
| Clipping x Competition | 1625 | 7 0.27 | 0.604 | 1) |
| Status x Fertilizer x Resource indicator value | 1636.53 | 0.00 | 0.878 | |
| Status x Fertilizer x Lifeform | 1643.06 | 0.45 | 0.730 | |
| Status x Fertilizer x Functional group | 1649.44 | 0.12 | 0.781 | |
| Status x Fertilizer x Competition | 1634.07 | 1.44 | 0.218 | |
| Status x Fertilizer x Clipping | 1638.81 | 0.01 | 0.888 | |
| **Status x Competition x Resource indicator value** | **1655.5** | **8.79** | **0.003** | |
| Status x Competition x Lifeform | 1638.76 | 1.95 | 0.238 | |
| Status x Competition x Functional group | 1618.54 | 1.53 | 0.222 | |
| Status x Clipping x Resource indicator value | 1654.13 | 0.48 | 0.658 | |
| Status x Clipping x Lifeform | 1644.61 | 0.29 | 0.706 | |
| Status x Clipping x Functional group | 1622.96 | 0.99 | 0.346 | |
| Status x Clipping x Competition | 1625.64 | 0.24 | 0.770 | |
| Fertilizer x Competition x Resource indicator value | 1634.63 | 0.10 | 0.606 | |
| Fertilizer x Competition x Lifeform | 1629.51 | 0.09 | 0.421 | |
| Fertilizer x Competition x Functional group | 1652.67 | 0.53 | 0.528 | |
| **Fertilizer x Clipping x Resource indicator value** | **1651.3** | **4.63** | **0.031** | |
| Fertilizer x Clipping x Lifeform | 1627.78 | 0.27 | 0.552 | |
| Fertilizer x Clipping x Functional group | 1647.76 | 0.32 | 0.572 | |
| **Fertilizer x Clipping x Competition** | **1651.6** | **4.86** | **0.027** | |
| Clipping x Competition x Resource indicator value | 1616.53 | 3.05 | 0.116 | |
| Clipping x Competition x Lifeform | 1631.42 | 0.04 | 0.915 | |
| Clipping x Competition x Functional group | 1623.97 | 0.33 | 0.564 | |
| Status x Fertilizer x Competition x RI | 1638.53 | 1.92 | 0.166 | |
| Status x Fertilizer x Competition x Lifeform | 1661.23 | 0.07 | 0.999 | |
| Status x Fertilizer x Competition x FG | 1659.33 | 0.11 | 0.227 | |
| Status x Fertilizer x Clipping x RI | 1663.16 | 0.02 | 0.885 | |
| Status x Fertilizer x Clipping x Lifeform | 1646.32 | 0.57 | 0.407 | |
| Status x Fertilizer x Clipping x FG | 1651.32 | 0.65 | 0.428 | |
| Status x Fertilizer x Clipping x Competition | 1640.8 | 0.90 | 0.436 | |
| Status x Clipping x Competition x RI | 1665.14 | 0.00 | 0.980 | |
| Status x Clipping x Competition x Lifeform | 1655.65 | 0.18 | 0.632 | |
| Status x Clipping x Competition x FG | 1627.41 | 3.05 | 0.100 | |
| Fertilizer x Clipping x Competition x RI | 1657.48 | 0.14 | 0.705 | |
| Fertilizer x Clipping x Competition x Lifeform | 1656.41 | 3.21 | 0.073 | |
| Fertilizer x Clipping x Competition x FG | 1667.13 | 0.00 | 0.883 | |
| ***Random terms*** | *Variance* | *SD* | | |
| Family | 0 | 0 | | |
| Group | 0.152 | 0.390 | | |
| Species | 0.209 | 0.457 | | |
| Block | 0.010 | 0.099 | | |

Table S3: Results from linear mixed effect models testing for the effects of Status, Fertilizer, experimental clipping and resource indicator value on the biomass of the competitor *L. perenne*, without (left) and with (right) taking the biomass of the target species into account. Significances were obtained by stepwise deletion of non-significant terms and comparing the resulting model to the previous ones using log-likelihood-ratio tests. This resulted in a minimal model containing only significant terms. We kept random factors in the model and present their variances. Number in bold indicate statistical significance. To obtain Ch$^2$ and *P*-values of two-way interactions and main effects, we excluded all three-way, respectively all higher-order interactions and compared this model with models missing the factors of interest (indicated by [1)]).

| **Fixed factors** | AIC | *Chi 2* | *P* | AIC | *Chi 2* | *P* |
|---|---|---|---|---|---|---|
| Biomass of target plants (log) | - | - | - | **200.94** | **28.0988** | **<0.0001** |
| Status | 200.85 | 4.9544 | 0.026 | 176 | 3.1662 | 0.0752 |
| Fertilizer | 215.88 | 19.9782 | <0.0001 | **195.44** | **22.6032** | **<0.0001** |
| Clipping | 218.13 | 19.1936 | <0.0001 [1)] | **184.39** | **11.555** | **0.0006** |
| Resource indicator value | 203.05 | 4.1163 | 0.042 [1)] | **180.07** | **7.2314** | **0.0071** |
| Functional group | 197.11 | 0.147 | 0.702 | 173.16 | 0.104 | 0.747 |
| Lifeform | 197.9 | 2.79 | 0.095 | 173.02 | 1.853 | 0.173 |
| Status x Resource indicator value | 206.15 | 0.032 | 0.858 | 180.32 | 0.05 | 0.823 |
| Status x Lifeform | 217.39 | 0.404 | 0.525 | 196.67 | 0.16 | 0.689 |
| Status x Functional group | 198.52 | 0.255 | 0.614 | 192.41 | 0.674 | 0.412 |
| Status x Fertilizer | 196.73 | 1.156 | 0.282 | 175.17 | 1.317 | 0.251 |
| Status x Clipping | 196.28 | 1.552 | 0.213 | 174.96 | 1.786 | 0.181 |
| Fertilizer x Resource indicator value | 201.19 | 0.211 | 0.646 | 176.67 | 0.121 | 0.728 |
| Fertilizer x Lifeform | 196.82 | 1.518 | 0.218 | 173.95 | 0.885 | 0.347 |
| Fertilizer x Functional group | 213.21 | 0.415 | 0.519 | 189.56 | 0.325 | 0.569 |
| Fertilizer x Clipping | 198.96 | 3.553 | 0.059 | 175.06 | 3.731 | 0.053 |
| Clipping x Resource indicator value | **200.94** | **5.0374** | **0.025** | 174.84 | 3.823 | 0.051 |
| Clipping x Lifeform | 197.41 | 2.593 | 0.107 | 173.33 | 1.381 | 0.24 |
| Clipping x Functional group | 197.57 | 1.054 | 0.305 | 175.86 | 1.187 | 0.276 |
| Status x Fertilizer x Resource indicator value | 208.11 | 0.003 | 0.954 | 182.27 | 0.048 | 0.826 |
| Status x Fertilizer x Lifeform | 218.99 | 0.251 | 0.616 | 198.51 | 0.198 | 0.656 |
| Status x Fertilizer x Functional group | 215.81 | 0.423 | 0.516 | 195.07 | 0.396 | 0.529 |
| Status x Fertilizer x Clipping | 204.2 | 0.05 | 0.824 | 184.22 | 0.024 | 0.878 |
| Status x Clipping x Resource indicator value | 210.11 | 0.003 | 0.958 | 186.2 | 0.006 | 0.941 |
| Status x Clipping x Lifeform | 222.56 | 0.027 | 0.87 | 200.31 | 0.08 | 0.778 |
| Status x Clipping x Functional group | 200.27 | 1.072 | 0.3 | 193.73 | 0.665 | 0.415 |
| Fertilizer x Clipping x Resource indicator value | 202.98 | 0.785 | 0.376 | 178.55 | 0.23 | 0.632 |
| Fertilizer x Clipping x Lifeform | 197.3 | 3.016 | 0.082 | 175.06 | 2.103 | 0.147 |
| Fertilizer x Clipping x Functional group | 214.79 | 0.978 | 0.323 | 191.24 | 0.829 | 0.362 |
| Status x Fertilizer x Clipping x Resource indicator value | 212.11 | 0.901 | 0.343 | 188.19 | 0.63 | 0.427 |
| Status x Fertilizer x Clipping x Lifeform | 224.54 | 0.104 | 0.747 | 202.23 | 0.353 | 0.553 |
| Status x Fertilizer x Clipping x FG | 220.74 | 0.175 | 0.676 | 203.88 | 0.006 | 0.936 |
| ***Random terms*** | *Variance* | *SD* | | *Variance* | *SD* | |
| Family | 0 | 0 | | 0 | 0 | |
| Group | 0.031 | 0.177 | | 0.023 | 0.152 | |
| Species | 0.015 | 0.123 | | 0.011 | 0.104 | |
| Block | 0.022 | 0.149 | | 0.026 | 0.160 | |

Table S4: Results from a linear mixed effect model testing for the effects of Status, Fertilizer, experimental clipping and resource indicator value on the percentage of ambient herbivore damage. Significances were obtained by stepwise deletion of non-significant terms and comparing the resulting model to the previous ones using log-likelihood-ratio tests. This resulted in a minimal model containing only significant terms. We kept random factors in the model and present their variances. Number in bold indicate statistical significance. To obtain $Ch^2$ and *P*-values of two-way interactions and main effects, we excluded all three-way, respectively all higher-order interactions and compared this model with models missing the factors of interest (indicated by [1)]).

| Fixed factors | AIC | *Chi 2* | *P* | |
|---|---|---|---|---|
| Status | -492.89 | 0.2065 | 0.650 | 1) |
| Fertilizer | -498.06 | 0.367 | 0.545 | |
| **Clipping** | **-487.04** | **6.0606** | **0.014** | 1) |
| Competition | -497.62 | 0.8076 | 0.369 | |
| Resource indicator value | -493.04 | 0.051 | 0.821 | 1) |
| Functional group (FG) | -496.43 | 3.796 | 0.051 | |
| Lifeform (LF) | -492.37 | 0.7271 | 0.394 | 1) |
| Status x Fertilizer | -494.17 | 0.351 | 0.554 | |
| Status x Clipping | -476.39 | 0.794 | 0.373 | |
| Fertilizer x Clipping | -472.4 | 0.253 | 0.615 | |
| Status x Competition | -483.75 | 0.003 | 0.960 | |
| Fertilizer x Competition | -486.56 | 0.323 | 0.570 | |
| Clipping x Competition | -498.22 | 3 | 0.083 | |
| **Status x Resource indicator value** | **-493.5** | **4.9328** | **0.026** | |
| Fertilizer x Resource indicator value | -489.5 | 0.161 | 0.688 | |
| Clipping x Resource indicator value | -497.27 | 0.01 | 0.919 | |
| Competition x Resource indicator value | -499.22 | 0.043 | 0.836 | |
| Status x Functional group | -475.19 | 0.147 | 0.701 | |
| Fertilizer x Functional group | -444.76 | 0.001 | 0.978 | |
| Clipping x Functional group | -477.23 | 1.161 | 0.281 | |
| Competition x Functional group | -460.77 | 0.564 | 0.453 | |
| Status x Lifeform | -492.52 | 0.176 | 0.675 | |
| Fertilizer x Lifeform | -495.02 | 1.155 | 0.282 | |
| **Clipping x Lifeform** | **-494.01** | **4.4145** | **0.036** | |
| Competition x Lifeform | -452.48 | 0.008 | 0.929 | |
| Status x Fertilizer x Resource indicator value | -487.66 | 0.902 | 0.342 | |
| Status x Fertilizer x Lifeform | -490.7 | 0.8 | 0.371 | |
| Status x Fertilizer x Functional group | -431.63 | 0.035 | 0.851 | |
| Status x Fertilizer x Competition | -480.06 | 0.001 | 0.979 | |
| Status x Fertilizer x Clipping | -466.64 | 0.09 | 0.764 | |
| Status x Competition x Resource indicator value | -481.75 | 0.307 | 0.579 | |
| Status x Competition x Lifeform | -450.49 | 0.048 | 0.827 | |
| Status x Competition x Functional group | -427.7 | 0.013 | 0.909 | |
| Status x Clipping x Resource indicator value | -459.34 | 0.372 | 0.542 | |
| Status x Clipping x Lifeform | -468.39 | 0.253 | 0.615 | |
| Status x Clipping x Functional group | -473.33 | 1.064 | 0.302 | |
| Status x Clipping x Competition | -456.02 | 0.08 | 0.777 | |
| Fertilizer x Competition x Resource indicator value | -484.89 | 0.865 | 0.352 | |
| Fertilizer x Competition x Lifeform | -448.54 | 0.023 | 0.879 | |
| Fertilizer x Competition x Functional group | -442.76 | 0.293 | 0.588 | |
| Fertilizer x Clipping x Resource indicator value | -463.55 | 0.521 | 0.471 | |
| Fertilizer x Clipping x Lifeform | -470.65 | 0.89 | 0.346 | |
| Fertilizer x Clipping x Functional group | -435.58 | 0.005 | 0.945 | |
| Fertilizer x Clipping x Competition | -469.54 | 0.847 | 0.357 | |
| Clipping x Competition x Resource indicator value | -495.28 | 1.742 | 0.187 | |
| Clipping x Competition x Lifeform | -439.33 | 0.203 | 0.652 | |
| Clipping x Competition x Functional group | -457.71 | 0.308 | 0.579 | |
| Status x Fertilizer x Competition x Resource indicator value | -478.06 | 1.171 | 0.279 | |
| Status x Fertilizer x Competition x Lifeform | -446.56 | 0.194 | 0.659 | |
| Status x Fertilizer x Competition x FG | -419.74 | 0.006 | 0.938 | |
| Status x Fertilizer x Clipping x Resource indicator value | -429.67 | 0.034 | 0.854 | |
| Status x Fertilizer x Clipping x Lifeform | -464.73 | 0.822 | 0.365 | |
| Status x Fertilizer x Clipping x FG | -425.72 | 0.011 | 0.916 | |
| Status x Fertilizer x Clipping x Competition | -441.05 | 0.276 | 0.599 | |
| Status x Clipping x Competition x Resource indicator value | -454.1 | 0.386 | 0.535 | |
| Status x Clipping x Competition x Lifeform | -423.73 | 0.007 | 0.932 | |
| Status x Clipping x Competition x FG | -421.73 | 0.007 | 0.933 | |
| Fertilizer x Clipping x Competition x Resource indicator value | -462.07 | 0.697 | 0.404 | |
| Fertilizer x Clipping x Competition x Lifeform | -437.53 | 0.052 | 0.819 | |
| Fertilizer x Clipping x Competition x FG | -433.59 | 0.047 | 0.828 | |
| ***Random terms*** | *Variance* | *SD* | | |
| Family | 0 | 0 | | |
| Group | 0.012 | 0.107 | | |
| Species | 0.005 | 0.069 | | |
| Block | 0.000 | 0.004 | | |

Fig. S1: Resource indicator values of common and rare plant species. High values indicate that a species originate from nutrient rich and moist habitats, low values indicate that a species originate from nutrient poor and dry habitats.

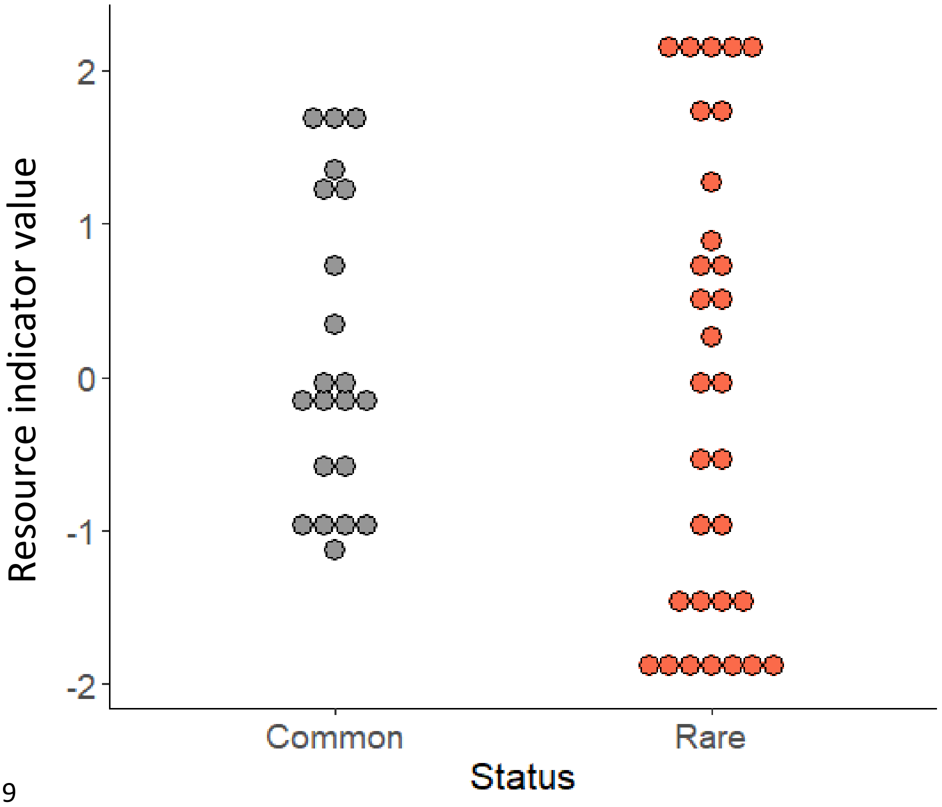

**Fig. S2:** Effects of competition (indicated by grass icon) and experimental clipping (indicated by a scissor) without and with fertilization (fertilized plants in brown) on aboveground biomass (back-transformed, in g). Shown are fitted estimates from a liner mixed effect model. Error bars indicate confidence intervals (obtained from the effect package in r). C= plant growing with competition, H = plant had been experimentally clipped, F = plants had been fertilized.

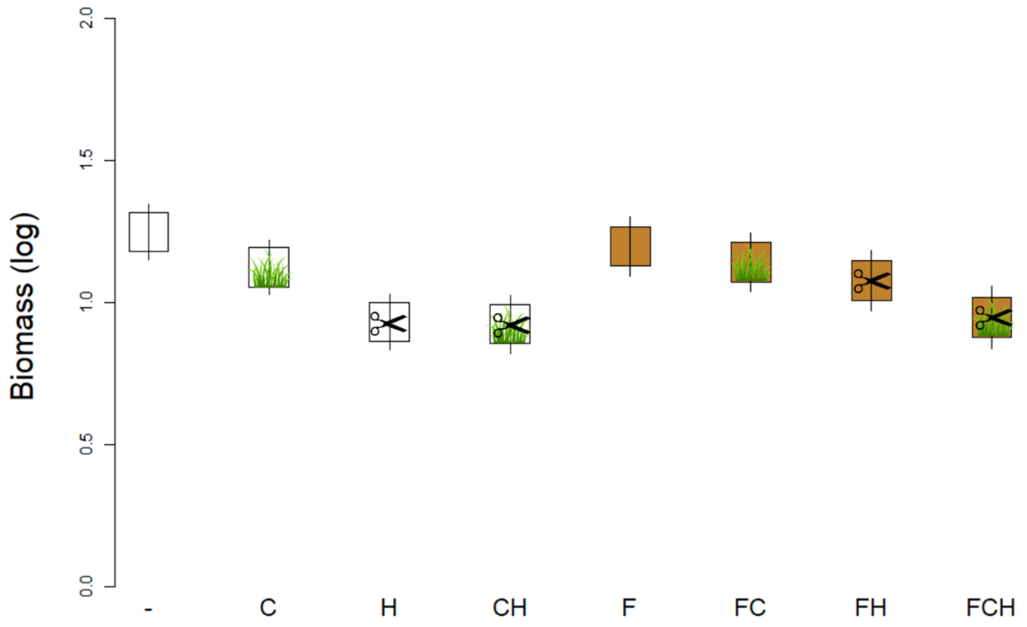

Figure S3: Effects of experimental clipping (indicated by a scissor) and fertilisation (fertilized plants in brown) on aboveground biomass (back-transformed, in g) of our plant species originating from low, medium or high resource habitats (resource indicator values -1.7, 0.15 and 2 respectively). Shown are fitted estimates from a liner mixed effect model. Error bars indicate confidence intervals (obtained from the effect package in r).

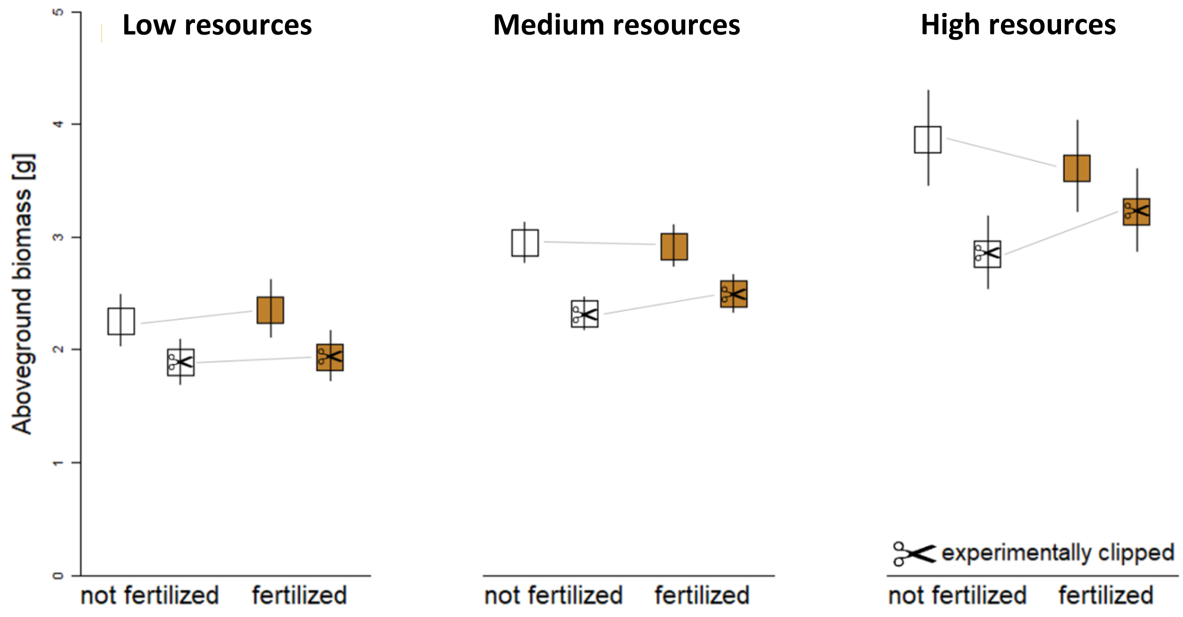
